# Biological invasions in rodent communities: from ecological interactions to zoonotic bacterial infection issues

**DOI:** 10.1101/108423

**Authors:** Christophe Diagne, M. Galan, Lucie Tamisier, Jonathan d’Ambrosio, Ambroise Dalecky, Khalilou Bâ, Mamadou Kane, Youssoupha Niang, Mamoudou Diallo, Aliou Sow, C. Tatard, A. Loiseau, O. Fossati-Gaschignard, Mbacké Sembène, Jean-François Cosson, Nathalie Charbonnel, Carine Brouat

## Abstract

Several hypotheses (such as ‘enemy release’, ‘novel weapon’, ‘spillback’ and ‘dilution/density effect’) suggest changes in host-parasite ecological interactions during biological invasion events. Such changes can impact both invasion process outcome and the dynamics of exotic and/or endemic zoonotic diseases. To evaluate these predictions, we investigated the ongoing invasions of the house mouse *Mus musculus domesticus*, and the black rat, *Rattus rattus*, in Senegal (West Africa). We focused on zoonotic bacterial communities depicted using 16S rRNA amplicon sequencing approach in both invasive and native rodents sampled along two well-defined invasion routes. Overall, this study provided new ecological evidence connecting parasitism and rodent invasion process, with diverse potential roles of zoonotic bacteria in the invasion success. Our results also highlighted the main factors that lie behind bacterial community structure in commensal rodents. Further experimental studies as well as comparative spatio-temporal surveys are necessary to decipher the actual role of zoonotic bacteria in these invasions. Our data also gave new support for the difficulty to predict the direction in which the relationship between biodiversity changes and disease risk could go. These results should be used as a basis for public health prevention services to design reservoir monitoring strategies based on multiple pathogen surveillance.

## Introduction

There is a close link between biological invasions (the establishment and spread of a non-native species in a new range) and parasites (here referred collectively to macroparasites and microparasites), with the potential for each to feed back on the other (Dunn and Hatcher 2015). First, parasites may alter the outcome of biological invasions by impacting host fitness and/or changing the strength of interactions between introduced and native hosts (Prenter, et al. 2004). As such, native parasites may prevent new host invasion if they infect introduced hosts and induce detrimental (direct and/or indirect) effects on their fitness (Strauss, et al. 2012). Also, native parasite pressure in the resident hosts may be reduced by introduced hosts through dilution or density effects (Keesing, et al. 2010), and indirectly contribute to increase the competitive ability of native hosts against invaders. Conversely, parasites may favour invasion success when introduction results in the loss of (a subset of) parasites by the exotic host (enemy release hypothesis; (Colautti, et al. 2004). Indeed the low number of introduced individuals may not carry the complete range of parasites found in source sites (founder effect) or introduced parasites may fail to thrive in the new environment (MacLeod, et al. 2010). Such parasite reduction is predicted to enhance the fitness and competitive ability of the introduced species in the new range. Moreover, parasites shared by both native and introduced hosts may also favour invasion success by inducing adverse effects on the fitness and/or survival of the native competitors. Invaders may introduce and transmit novel parasites to native hosts (“novel weapon” hypothesis; (Strauss, et al. 2012) or they may acquire and better amplify endemic parasites, resulting in higher prevalence/abundance in native hosts than expected in the absence of the new hosts (spillback hypothesis; (Kelly, et al. 2009).

Second, invasion related changes in host community structure and diversity may *in fine* influence the dynamics of diseases potentially affecting wildlife, livestock and/or human health (Young, et al. 2017). Disease emergence events associated with non-native pathogens brought by animal invaders have already been observed (Dunn and Hatcher 2015). Despite a growing interest these last decades, there are still gaps in the basic studies devoted to relationships between parasite ecology, disease dynamics and invasion process (Young, et al. 2017). Notably, few studies have examined the concomitant effects of both native and invasive host communities on infectious disease risks in invaded ecosystems.

In this paper, we examined ecological interactions between native and invasive host rodents and their bacterial communities in the context of well-studied ongoing invasions in Senegal (West Africa). They concern two major invasive species (Global Invasive Species Database - http://www.issg.org/database/): the house mouse *Mus musculus domesticus* (Dalecky, et al. 2015) Lippens et al. unpublished data) and the black rat *Rattus rattus* (Konecny, et al. 2013). Historical records and molecular analyses have shown that these rodents were first brought to Senegalese coasts by Europeans at colonial times, and remained in coastal villages and towns until the beginning of the 20^th^ century. Both taxa have spread further inland during the last century via well-defined invasion routes, due to the increase of human activities and the improvement transport infrastructures in mainland Senegal. The distribution of the house mouse covers now most of the northern and central part of Senegal, while the black rat is distributed in the southern part of the country. Along their invasion routes, the invasive species have colonised the human settlements (either towns or small villages) from which they have progressively weed out the commensal native rodent species, the Guinea multimammate mouse *Mastomys erythroleucus* and the natal multimammate mouse *Mastomys natalensis*. At invasion fronts, both invasive and native species cohabit (Dalecky, et al. 2015; Granjon and Duplantier 2009). The role of parasites in successful colonization events of the house mouse and black rat has previously been documented, mainly with regard to insular ecosystems (Harris 2009; Wyatt, et al. 2008). More recently, (Diagne, et al. 2016) have revealed an association between the loss of gastrointestinal helminths and the range expansion of these rodents in Senegal. Because invasive and African rodent species studied here are known to carry a large array of zoonotic agents (Meerburg, et al. 2009), it is also likely that the risks of zoonotic diseases for human may change along the mouse and rat invasion routes.

Here, we focused on the zoonotic bacteria detected in rodents using the 16S rRNA amplicon sequencing approach described in (Galan, et al. 2016). This community comprises several pathogens that may be transmitted to other mammal species including humans (Han, et al. 2015; Meerburg, et al. 2009). We first analysed the relationships between rodent-bacteria interactions and ongoing invasions by testing for patterns expected under the ecological processes described above: (i) a decrease of bacterial species richness and prevalence levels in invasive rodents at their invasion fronts compared to their original coastal sites, as expected under the enemy release hypothesis, (ii) changes in bacterial species richness and prevalence levels in native rodents from invaded areas compared to non-invaded areas as expected under the novel weapon/spillback (increase expected for bacteria shared by native and invasive rodents at invasion fronts) or dilution/density effect (decrease expected) hypotheses. We next discussed the importance of these variations in bacterial communities observed along rodent invasion routes in terms of public health.

## Results

A total of 985 rodents were sampled in 24 sites. Trapping sessions showed variations in rodent communities along both invasion routes, from western coastal sites of long-established invasion to inland invasion fronts and non-invaded sites (Table 1; Fig. 1).

**Table 1:** Sampling sites and their invasion status as well as rodent sample sizes and proportion of infected hosts in each site on a) the mouse invasion route and b) the rat invasion route. Legend: LI: sites of long-established invasion; IF: recently invaded sites (invasion front); NI: non-invaded sites; Lat: latitude of sites; Long: longitude of sites; N: total number of rodents sampled; M: males; F: females; n: number of hosts screened for bacteria; P: prevalence (number of infected hosts / number of screened hosts) in %. ‘-’ indicates no data (no rodent sampled or screened).

| Sites (Invasion status) | Lat (North) | Long (West) | *Mus musculus domesticus* |  | *Mastomys erythroleucus* |  |
| --- | --- | --- | --- | --- | --- | --- |
|  |  |  | N (M/F) | n (P) | N (M/F) | n (P) |
| Dagathie (LI) | 15.62 | -16.25 | 30 (18/12) | 24 (66.7) | - | - |
| Mbakhana (LI) | 16.08 | -16.36 | 30 (14/16) | 26 (19.2) | - | - |
| Ndombo (LI) | 16.43 | -15.70 | 23 (10/13) | 21 (38.4) | - | - |
| Thilene (LI) | 16.26 | -16.18 | 26 (9/17) | 19 (47.4) | - | - |
| Aere Lao (IF) | 16.40 | -14.32 | 50 (22/28) | 25 (60) | 22 (13/9) | 20 (60) |
| Croisement Boube (IF) | 16.49 | -15.00 | 45 (24/21) | 27 (18.5) | 3 (2/1) | - |
| Dendoudi (IF) | 15.39 | -13.53 | 19 (13/5) | 15 (33.3) | 28 (18/10) | 22 (91) |
| Dodel (IF) | 16.48 | -14.42 | 24 (9/15) | 20 (45) | 33 (16/17) | 20 (30) |
| Lougue (IF) | 16.06 | -13.91 | 28 (13/15) | 25 (28) | 25 (15/10) | 20 (65) |
| Diomandiou Walo (NI) | 16.49 | -15.00 | - | - | 14 (10/4) | 13 (84.6) |
| Doumnga Lao (NI) | 16.33 | -14.22 | - | - | 43 (18/25) | 20 (35) |
| Lambango (NI) | 15.46 | -13.53 | - | - | 23 (10/13) | 21 (85.7) |
| Sare Maounde (NI) | 15.98 | -13.91 | - | - | 9 (5/4) | 9 (33.3) |

b)
| Sites (Invasion status) | Lat (North) | Long (West) | *Rattus rattus* |  | *Mastomys erythroleucus* |  | *Mastomys natalensis* |  |
| --- | --- | --- | --- | --- | --- | --- | --- | --- |
|  |  |  | N (M/F) | n (P) | N (M/F) | n (P) | N (M/F) | n (P) |
| Diakene-Wolof (LI) | 12.45 | -16.64 | 30 (12/18) | 24 (25) | 8 (5/3) | - | - | - |
| Diattacounda (LI) | 12.57 | -15.68 | 35 (19/16) | 27 (70.4) | 16 (9/7) | - | - | - |
| Marsassoum (LI) | 12.83 | -15.97 | 29 (13/16) | 25 (76) | 3 (0/3) | - | - | - |
| Tobor (LI) | 12.66 | -16.25 | 29 (12/17) | 21 (81) | 1 (1/0) | - | - | - |
| Badi Nieriko (IF) | 13.37 | -13.37 | 73 (35/38) | 21 (62) | 14 (6/8) | 13 (84.6) | - | - |
| Boutougoufara (IF) | 13.39 | -12.48 | 40 (15/25) | 28 (28.6) | 21 (12/9) | 21 (95.2) | - | - |
| Kedougou (IF) | 12.55 | -12.17 | 34 (13/21) | 21 (9.5) | - | - | 30 (11/19) | 22 (72.7) |
| Soutouta (IF) | 13.80 | -12.71 | 26 (13/13) | 22 (86.4) | 12 (5/7) | 12 (75) | - | - |
| Bransan (NI) | 13.26 | -12.10 | - | - | 6 (4/2) | - | 33 (14/19) | 23 (78.3) |
| Mako (NI) | 12.85 | -12.35 | - | - | - | - | 48 (28/20) | 24 (95.8) |
| Segou (NI) | 12.41 | -12.28 | - | - | - | - | 26 (10/16) | 24 (100) |

**Figure 1:**
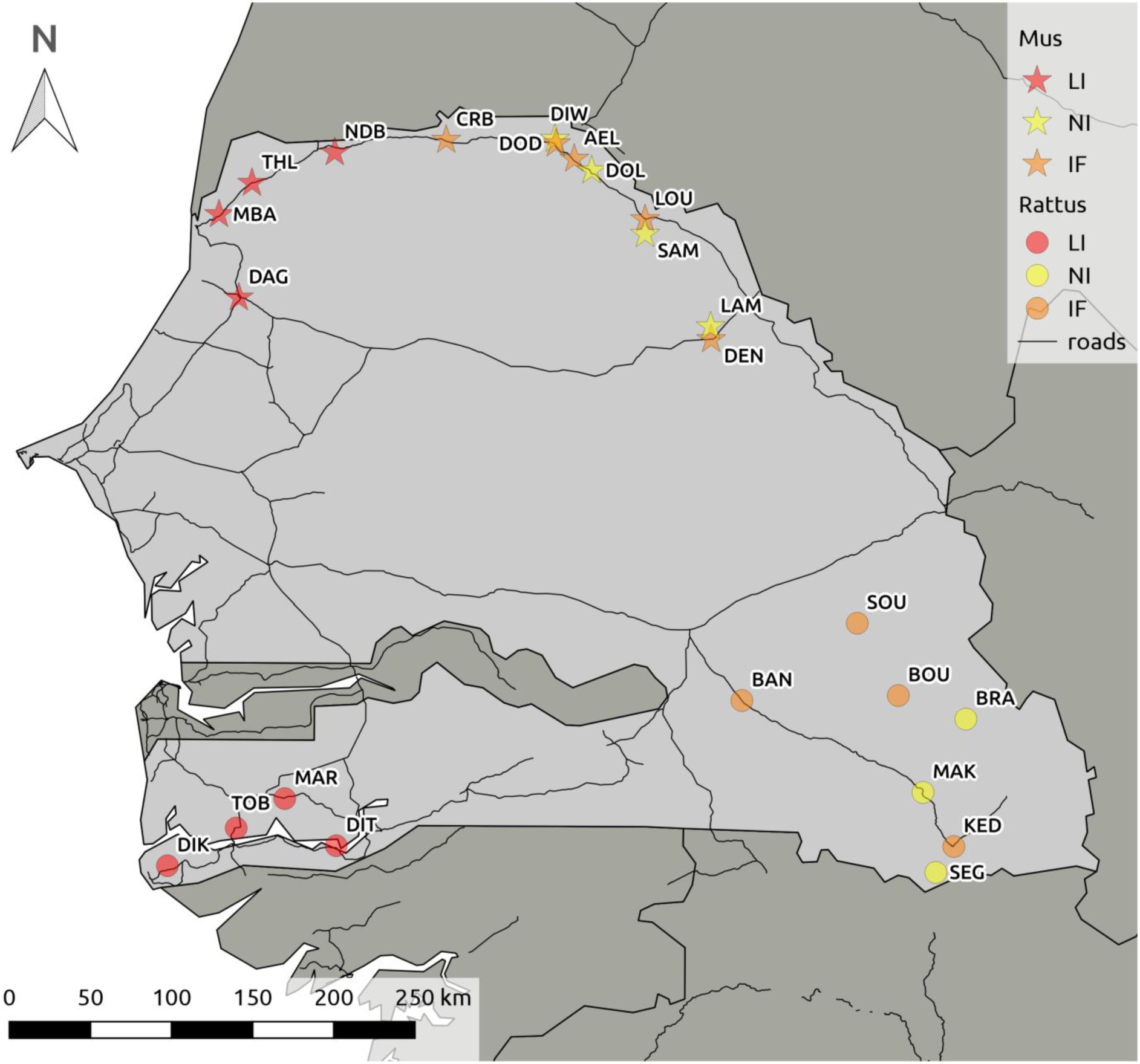
Rodent sampling sites on the house mouse (*Mus musculus domesticus*) (star symbols) and black rat (*Rattus rattus*) (circle symbols) invasion routes in Senegal (West Africa). Colour legend: red for sites of long-established invasion; orange for recently invaded sites; yellow for non-invaded sites. Correspondence between site codes and names are detailed in Table 1.

Along the mouse invasion route (Table 1a), only invasive *M. m. domesticus* were captured in sites of long-established invasion. At the invasion front, native *Ma. erythroleucus* co-occurred generally with *M. m. domesticus*, although being nearly systematically less abundant. The only exception was the site of Croisement Boubé, where *Ma. erythroleucus* was not detected despite the recent introduction of *M. m. domesticus* in this area (Dalecky et al. 2015). As expected, only native rodents were sampled in non-invaded sites.

Along the rat invasion route (Table 1b), native rodent communities belonged to two sister species, *Ma. erythroleucus* and *Ma. natalensis*. Sites of long-established invasion were highly dominated by invasive *R. rattus*. At the invasion front, invasive *R. rattus* co-occurred with *Ma. erythroleucus* in three of the four sites, and with *Ma. natalensis* in the southernmost site (Kedougou). Only native rodents were sampled in non-invaded sites, and they mainly belonged to *Ma. natalensis*. Some sparse *Ma. erythroleucus* individuals were sampled in one non-invaded site (Bransan) and in sites of long-established invasion (cf. Table 1), probably coming from the surrounding fields where this rodent species is generally abundant (Granjon and Duplantier 2009). They were not included in further statistical analyses due to low sample sizes.

Bacteria were screened in 679 rodents (Table 1). Twelve zoonotic bacterial Operational Taxonomic Units (OTUs) were described (Galan et al. 2016): *Bartonella*, *Borrelia*, *Ehrlichia*, *Mycoplasma_1*, *Mycoplasma_2*, *Mycoplasma_3*, *Mycoplasma_4*, *Mycoplasma_5*, *Mycoplasma_6*, *Orientia*, *Rickettsia* and *Streptobacillus*.

**Distribution of bacterial communities along invasion routes.** Along the mouse invasion route, six zoonotic OTUs were found (Table 2): *Bartonella*, *Borrelia*, *Ehrlichia*, *Mycoplasma_1*, *Mycoplasma_3* and *Orientia*. *Orientia* was found exclusively in *M. m. domesticus*. Otherwise, all the OTUs were found in both *Ma. erythroleucus* and *M. m. domesticus*. *Mycoplasma_1*, *Ehrlichia* and *Borrelia* were found in all categories of sites (sites of long-established invasion, invasion front and non invaded sites). *Bartonella* and *Mycoplasma_3* were detected in *M. m. domesticus* in only one site of long-established invasion, while these OTUs were found in *Ma. erythroleucus* in both invasion front and non-invaded sites.

**Table 2:**
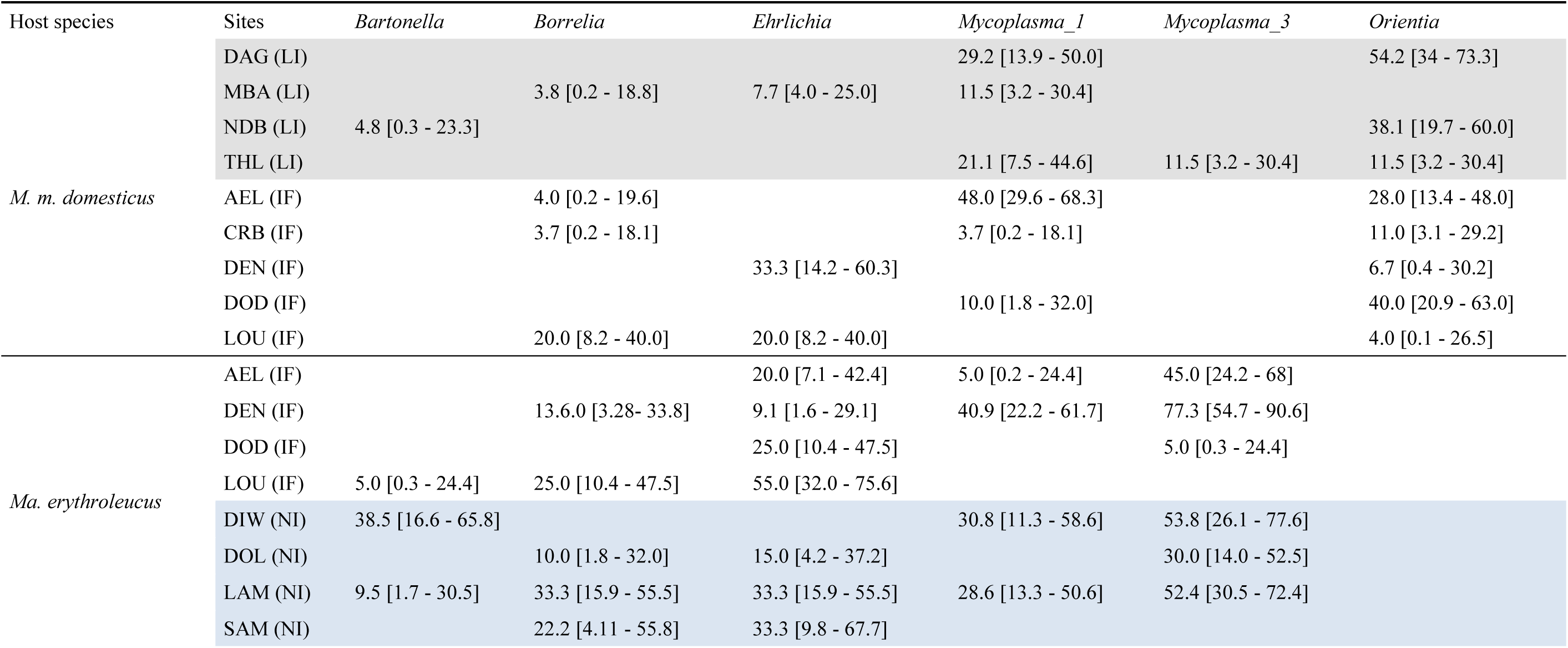
Prevalence in % [with 95% confidence intervals calculated with Sterne’s exact method] of bacterial OTUs detected in *Mus musculus domesticus* and *Mastomys erythroleucus* for each site sampled on the house mouse invasion route. LI: site of long-established invasion (only invasive host species); IF: invasion front (invasive + native host species); NI: non-invaded sites (only native host species). AEL: Aere Lao; CRB: Croisement Boube; DAG: Dagathie; DEN: Dendoudi; DIW: Diomandou Walo; DOD: Dodel; DOL: Doumnga Lao; LAM: Lambago; LOU: Lougue; MBA: Mbakhana; NDB: Ndombo; THW: Thiewle; THL: Thilene.

Along the rat invasion route, twelve pathogenic OTUs were recorded (Table 3): *Bartonella, Borrelia, Ehrlichia, Mycoplasma_1, Mycoplasma_2, Mycoplasma_3, Mycoplasma_4, Mycoplasma_5, Mycoplasma_6, Orientia, Rickettsia* and *Streptobacillus*. Four OTUs showed host specificity: *Mycoplasma_2, Mycoplasma_4, Rickettsia* in *R. rattus, Orientia* in *Ma. natalensis, Mycoplasma_5* and *Mycoplasma_6* in *Mastomys* spp. The six OTUs detected in both native and invasive rodents exhibited different distribution patterns. *Bartonella* was found in all categories of sites. *Borrelia* and *Ehrlichia* were restricted to one site at the invasion front. *Mycoplasma_1* and *Mycoplasma_3* were detected in *R. rattus* in one site only at the invasion front and in *Mastomys* spp. in both invasion front and non-invaded sites.

**Table 3:**
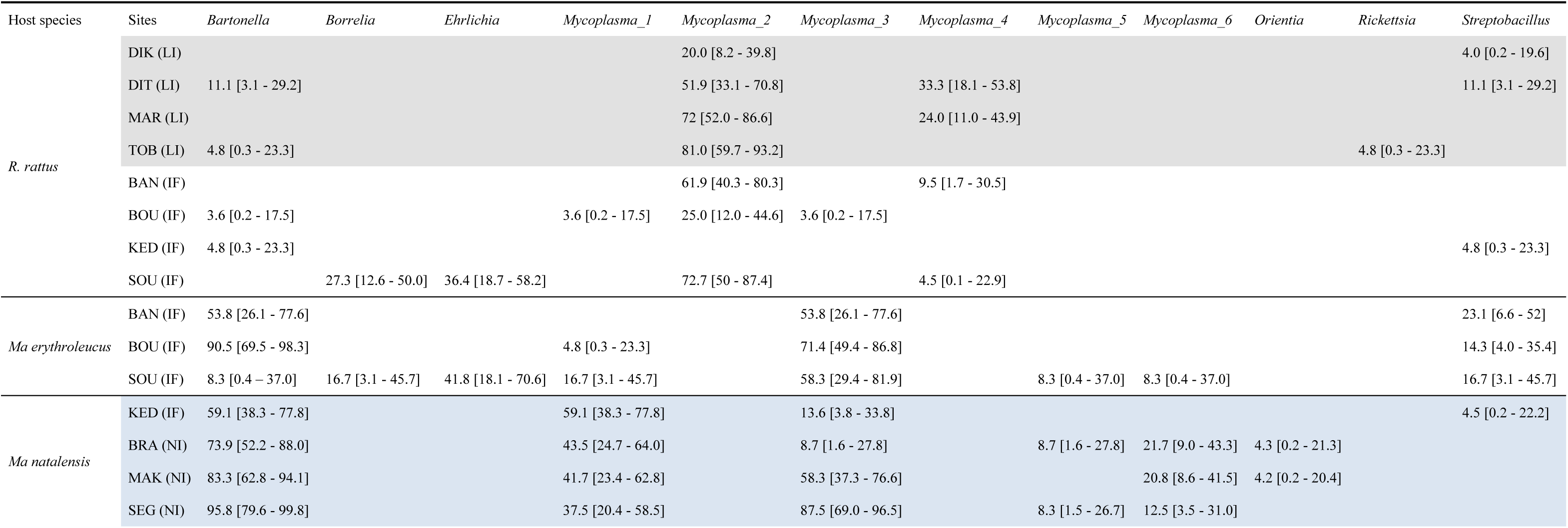
Prevalence in % [with 95% confidence intervals calculated with Sterne’s exact method] of bacterial OTUs detected in *Rattus rattus*, *Mastomys erythroleucus* and *Mastomys natalensis* for each sampling site from the black rat invasion route (details on codes for sites are provided in table 1). LI: site of long-established invasion; IF: invasion front (invasive + native host species); NI: non-invaded sites (only native host species). BAN: Badi Nieriko; BOU: Boutougoufara; BRA: Bransan; DIT: Diattacounda; DIK: Diakene-Wolof; KED: Kedougou; MAK: Mako; MAR: Marsassoum; SEG: Segou; SOU: Soutouta; TOB: Tobor.

For both invasion routes, hierarchical clustering and permutational multivariate analysis of variance (permanova) carried out on Bray-Curtis (BC) dissimilarity index-based matrices revealed that the bacterial communities differed significantly between invasive and native host species (mouse invasion route: BC percentage of total variance explained R2 = 0.29; F = 5.77, *p = 0.002*, Fig. 2; rat invasion route: BC R2 = 0.52; F = 14.35, *p < 0.001*, Fig. 3), but not between invasion-related categories of sites at intraspecific level (mouse invasion route: BC R2 = 0.05; F = 0.53, *p = 0.817*, Fig. 2; rat invasion route: BC R2 = 0.08; F = 1.13, *p = 0.334*, Fig. 3). The same analyses performed using the presence/absence data-based Jaccard dissimilarity index showed similar bacterial community structure patterns (Supplementary materials, Fig. 1, Fig. 2).

**Figure 2:**
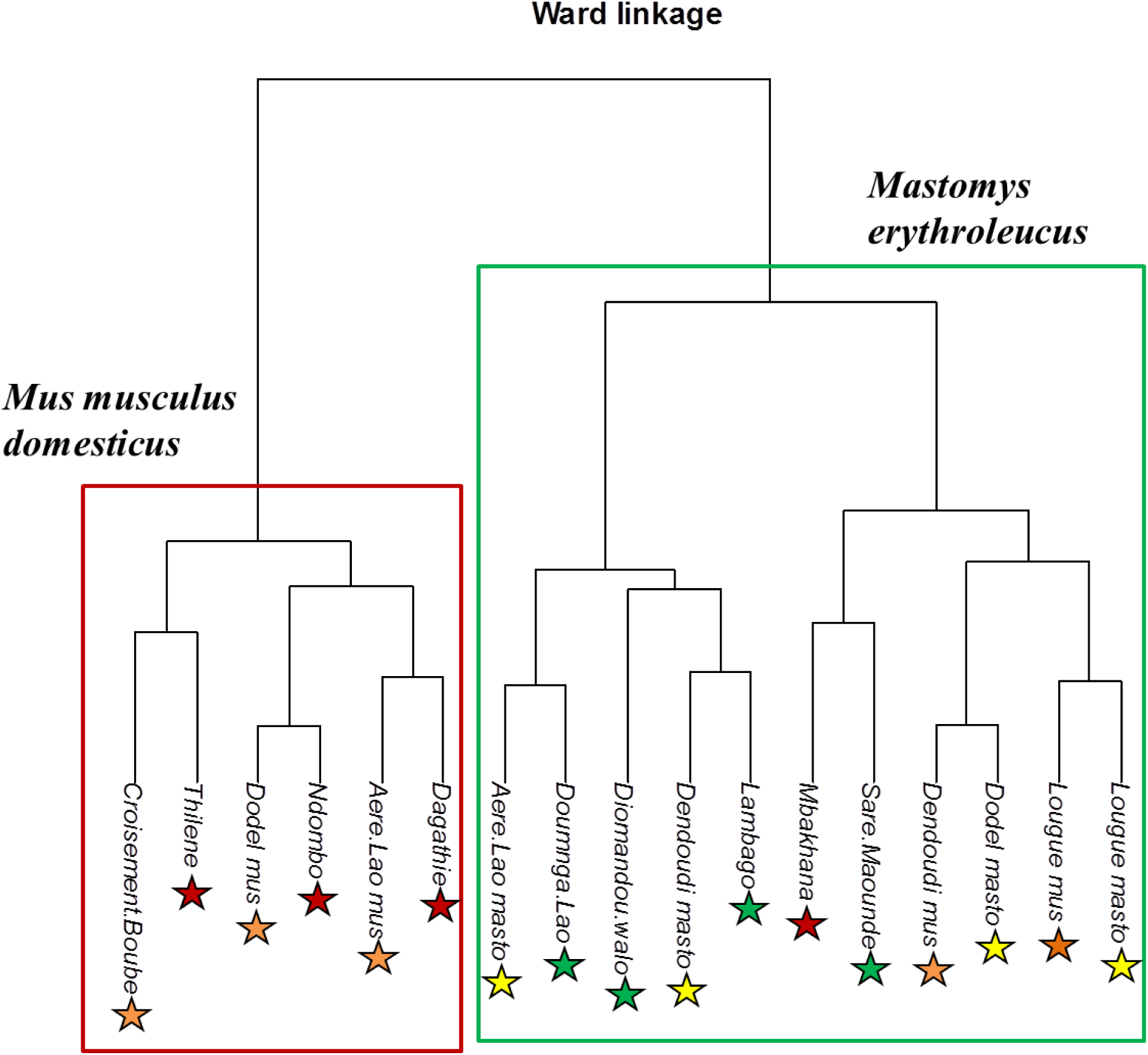
Bray-Curtis (BC) dissimilarity index-based Ward’s hierarchical clustering of the bacterial communities described in the rodent host populations sampled along the mouse invasion route. The graph shows that the bacterial communities were clustered by the host species (permutational multivariate analyses of variance [permanova] performed on BC index-based matrices: percentage of total variance explained R2 = 0.29; F = 5.77, *p = 0.002*) but not by the invasion status of sampling sites (permanova performed on BC index-based matrices: R2 = 0.05; F = 0.53, *p = 0.817*). Legend: red stars: *Mus musculus domesticus* populations at sites of long-established invasion; orange stars: *M. m. domesticus* populations at invasion front sites; yellow stars: *Mastomys erythroleucus* populations at invasion front sites; green stars: *Ma. erythroleucus* populations at non-invaded sites. At invasion front sites where both native and invasive species co-occurred, “mus” or “masto” was added after the site name for distinguishing *M. m. domesticus* and *Ma. erythroleucus* populations, respectively.

**Figure 3:**
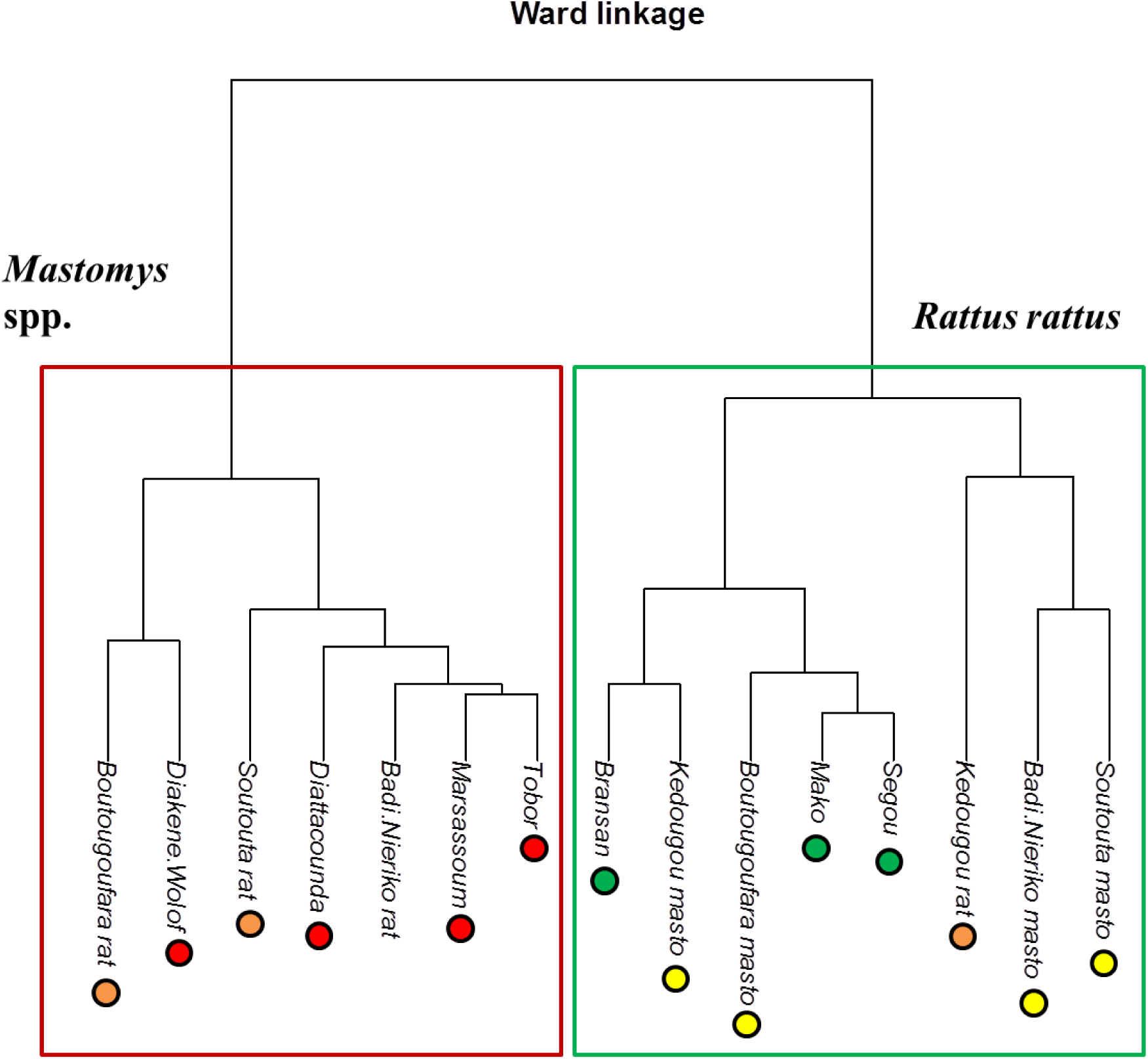
Bray-Curtis (BC) dissimilarity index-based Ward’s hierarchical clustering of the bacterial communities described in the rodent host populations sampled along the rat invasion route. The graph shows that the bacterial communities were clustered by the host species (permutational multivariate analyses of variance [permanova] performed on BC index-based matrices: percentage of total variance explained R2 = 0.52; F = 14.35, *p < 0.001*) but not by the invasion status of sampling sites (permanova performed on BC index-based matrices: R2 = 0.08; F = 1.13, *p = 0.334*). Legend: red stars: *Rattus rattus* populations at sites of long-established invasion; orange stars: *R. rattus* populations at invasion front sites; yellow stars: *Mastomys* spp. populations at invasion front sites; green stars: *Mastomys* spp. populations at non-invaded sites. At invasion front sites where both native and invasive species co-occurred, “rat” or “masto” was added after the site name for distinguishing *R. rattus* and *Mastomys* spp. populations, respectively.

**Factors shaping bacterial communities.** Generalized Linear Mixed Models (GLMMs) were used to decipher the effects of the site invasion status (e.g., site of long-established invasion, invasion front or non-invaded site) and of individual factors (gender, body mass of the rodent) on individual OTU species richness (number of OTUs found in a single host individual) or OTUs prevalence (proportion of hosts infected with a given OTU).

***Bacterial OTUs richness.*** Considering the mouse invasion route, a significant effect of *invasion status* (*p = 0.024*) indicated less bacterial OTUs in *M. m. domesticus* than in *Ma. erythroleucus* (pairwise Wilcoxon tests with Holm’s correction (WH tests), *p < 0.05*). There was no difference of species richness between *M. m. domesticus* from invasion front and from long-established invasion sites (WH test, *p = 0.99*), or between *Ma. erythroleucus* from invasion front and from non-invaded sites (WH test, *p = 0.39*). Species richness was higher in males than in females (*gender* effect: *p = 0.001*).

Considering the rat invasion route, *invasion status* (*p < 0.001*) had also a significant effect on species richness. Invasive *R. rattus* had less bacterial OTUs than native *Mastomys* rodents (WH tests, *p < 0.05*), and *Mastomys* rodents had less bacterial OTUs at the invasion front than in non-invaded sites (WH test, *p = 0.008*). No significant difference was detected between *R. rattus* populations from invasion front and long-established invasion sites (WH test, *p = 0.580*). Finally, male and heavier rodents harboured more OTUs than female and lighter ones (*p < 0.001* for *gender* and *body mass* effects).

***Bacterial OTUs prevalence.*** Considering the mouse invasion route, *invasion status* (*p < 0.001*) was significant for *Ehrlichia* only, indicating higher infection levels in *Ma erythroleucus* at the invasion front (WH tests, *p < 0.005*). Besides, males were more infected by *Mycoplasma_1* (*p < 0.001*), *Ehrlichia* (*p = 0.011*) and *Borrelia* (*p = 0.002*) than females. A significant interaction was found between *gender* and *invasion status* (*p = 0.014*) for *Mycoplasma_3*, suggesting that males were more infected in non-invaded sites than at invasion front, and the opposite trend for females. Finally, heavier rodents were less infected by *Ehrlichia* than lighter ones (*p = 0.039*).

Considering the rat invasion route, *invasion status* was significant for *Bartonella* (*p < 0.001*) and *Mycoplasma_6* (*p = 0.006*). At invasion front, *R. rattus* were less infected by *Bartonella* than *Mastomys* rodents (WH test, *p < 0.001*). *Mastomys* rodents were also less infected by *Bartonella* (WH test, *p = 0.001*) and *Mycoplasma_6* (WH test, *p = 0.004*) at the invasion front than in non-invaded sites. The factor *gender* was significant for *Bartonella* (*p = 0.025*), *Mycoplasma_1* (*p = 0.004*), *Mycoplasma_4* (*p = 0.02*), and *Mycoplasma_6* (*p = 0.003*) with higher infection levels observed in males than in females. The factor *body mass* significantly explained the prevalence of *Mycoplasma 1* (*p < 0.001*), *Mycoplasma_2* (*p = 0.001*), *Mycoplasma 4* (*p = 0.003*) and *Mycoplasma_5* (*p = 0.036*), with heavier rodents being more infected than lighter ones.

## Discussion

In this study, we examined variations of bacterial communities in invasive and native rodents sampled along two invasion routes in Senegal. Our results have several implications regarding (i) the factors that lie behind bacterial community structure, (ii) the ecological hypotheses relating parasitism to invasion outcome in commensal rodents, and (iii) the putative relationships between host biodiversity and disease risks.

Our study first confirmed that host individual factors such as body mass (Moore and Wilson 2002) and gender (Zuk and Stoehr 2010) are important drivers of bacterial infections in small mammals. In accordance with previous works (but see (Kiffner, et al. 2013)), we found that larger and male rodents were significantly more infected than lighter and female ones. Male-biased parasitism in rodents could be due to profound sexual differences in anatomy, physiology, behaviour, and evolutionary roles that could increase probabilities of both encountering and acquiring parasites (reviewed in (Krasnov, et al. 2012). Larger individuals may have greater home range, what may increase contact rates with parasites (Boyer, et al. 2010). Besides, as body mass can be considered as a proxy of host age, the positive correlation that is generally found between infection and body mass could correspond to higher levels of infection in older individuals. This pattern is likely to result from a longer time of exposure in older than younger rodents. Alternatively, the negative relationship found between *Ehrlichia* infection and body mass on the mouse invasion route may suggest that older individuals would be less susceptible to the infection, or that infected individuals died rapidly.

Establishing the origin of parasites (either native or exotic) is a key element in the context of the hypotheses relating parasitism and invasion success. The distribution of bacterial OTUs among native and invasive rodents may give useful insights when investigating their origin. As already found in rodents (Badenhorst, et al. 2012), some of the OTUs recorded here appeared to be relatively host specific. Indeed, three OTUs (*Mycoplasma_3*, *Mycoplasma_5*, *Mycoplasma_6*) were mainly found in *Mastomys* rodents, and are thus probably native. This could also be the case for *Ehrlichia,* which was detected at high prevalence in native rodents, and almost exclusively at invasion fronts in invasive rodents. The *Borrelia* OTU found in native and invasive rodents is also probably native, as large surveys have detected only one local species (e.g., *Borrelia crocidurae*) in small mammal communities of West Africa (Trape, et al. 2013; Vial, et al. 2006). Conversely, two OTUs (*Mycoplasma_2*, *Mycoplasma_4* along the rat invasion route) were found repeatedly and exclusively in invasive *R. rattus*. The distribution of these bacteria suggested that they have been introduced in Senegal by exotic *R. rattus*. A high susceptibility of *Mastomys* species to these OTUs may prevent the detection of infected native rodents. The detection of *Streptobacillus* in *R. rattus* at long-established invasion sites and in native rodents only at the invasion front of *R. rattus*, also indicated an exotic origin for this OTU. In a previous work, we suggested an exotic origin for *Orientia*, which is found exclusively in *M. m. domesticus* along the mouse invasion route (Cosson, et al. 2015). However, the detection of identical sequences of this OTU in some native *Ma. natalensis* samples from outside the range of *M. m. domesticus* on the otherside of Senegal raises questions about the exotic origin of this bacterial OTU or the involvement of host species other than exotic rodents in its introduction (e.g., migratory birds: (Varma 1964)).

As expected, bacterial communities varied along invasion routes and between invasive and native rodents. A main result was that bacterial species richness was globally higher in native rodents than in invasive ones. This pattern may reflect a loss of zoonotic bacteria experienced by *M. m. domesticus* and *R. rattus* at the time of their introduction in Senegal. Signatures of bacterial loss were however not observed for potentially exotic OTUs specifically carried by *M. m. domesticus* (i.e., *Orientia*) and *R. rattus* (i.e., *Mycoplasma_2* and *Mycoplasma_4*) along their respective invasion routes. In contrast to previous surveys of gastrointestinal (Diagne, et al. 2016), there was thus no support for a bacterial release hypothesis favouring the spread of *M. m. domesticus* and *R. rattus* in Senegal. Possible explanations could rely on the mode of transmission and/or vector specificity of the introduced bacteria. All bacterial OTUs mentioned here (except *Mycoplasma* and *Streptobacillus*) are vectored by arthropods (e.g., fleas, ticks, see (Galan, et al. 2016) for a synthesis). For instance, trombiculid mites involved in the transmission of *Orientia* (Paris, et al. 2013) are known for their very large host spectrum and low host specificity (Shatrov and Kudryashova 2006), which may have facilitated the introduction of the bacteria into new environments. The direct cycle of non-vectored bacteria such as *Mycoplasma* may also enhance their establishment probabilities.

Data on native rodents provided evidence in favour of ecological processes connecting parasitism and invasion success. The higher prevalence of *Ehrlichia* in native rodents at the mouse invasion front compared to non-invaded sites could result from a spillback process. The ‘novel weapon’ hypothesis might explain the relatively high prevalence of *Streptobacillus* in native *Mastomys* spp. at the invasion front of *R. rattus*. Indeed, this OTU is known to be common and commensal in rats, but putatively pathogenic in other rodents (Elliott 2007). Finally, our data also revealed patterns that are consistent with the dilution/density hypotheses on the rat invasion route. Indeed, species richness as well as *Bartonella* and *Mycoplasma_6* prevalence in native rodents were shown to be lower at invasion front than in non-invaded areas. The non-detection of *Mycoplasma_6* in *R. rattus* did not support an involvement of the exotic rodent as a sink reservoir, and may thus suggest a host density (i.e., decrease of native host densities due to the presence of invaders (Keesing, et al. 2010)) rather than a host dilution effect. In contrast, the occurrence *Bartonella* in *R. rattus* may suggest a dilution effect by the exotic rodent. Dilution effects were already shown by longitudinal surveys for *Bartonella* infections in native woodmouse *Apodemus sylvaticus* communities invaded by the bank vole *Myodes glareolus* in the United Kingdom (Telfer, et al. 2007).

Overall, this study evidenced diverse potential roles of zoonotic bacteria in the invasion success of rodents in Senegal. Because the ecological patterns described here did not enable to measure the impact of zoonotic bacteria at both host individual and population levels, experimental studies are necessary to decipher the role of zoonotic bacteria in the invasion process. Besides, developing comparative spatio-temporal surveys of various invasion routes is a further critical step to understand if it is possible to generalise on the relationships between parasitism and invasion success, despite environmental and demographic stochasticity (Colautti and Lau 2015).

Research on bacterial communities infecting wild rodents remains scarce in continental Africa. To our knowledge, it mainly concerned *Yersinia pestis* (e.g., (Moore, et al. 2015)), *Borrelia* sp. (Trape, et al. 2013), *Bartonella* sp. (Hayman, et al. 2013), and *Leptospira* sp. (Dobigny, et al. 2015). Our work allowed to identify four other genera of zoonotic bacteria (e.g., *Ehrlichia*, *Orientia*, *Rickettsia*, *Streptobacillus*) in invasive and/or native commensal rodents from Senegal, which may cause severe human diseases still underdiagnosed in Africa (WHO 2009). By doing so, this study corroborated previous works highlighting the major role of exotic rodents in public health issues (e.g., (Himsworth, et al. 2013). Indeed, we showed that invasive rodents have probably brought exotic bacteria at invasion fronts (e.g. *Orientia* in North Senegal by *M. m. domesticus*; *Streptobacillus* or *Rickettsia* in South-East Senegal by *R. rattus*). We also showed that the introduction of invasive rodents may change the prevalence of some bacteria carried by native rodents (e.g., increase of infections related to *Ehrlichia*; decrease of infections related to *Bartonella*), or lead to the replacement of local lineages or species of *Mycoplasma* by exotic ones. These results advocated for an increased vigilance along the invasion roads of these exotic rodents with regard to the emergence of rodent borne diseases.

In conclusion, our focus on invasion and parasitism provided a larger support to the assumption still strongly debated (Salkeld, et al. 2015) that the relationship between biodiversity and disease risk is difficult to predict, especially when tracking multiple diseases (Wood, et al. 2014). We advocate for public health prevention strategies based on multiple pathogen surveillance, especially in the case of invasive reservoir/vector species monitoring (Caffrey, et al. 2014). This is particularly true for underdeveloped regions, where many infectious diseases originating from wildlife are underreported.

## Methods

**Sample collection.** We used data from historical records and population genetics studies (Dalecky, et al. 2015; Konecny, et al. 2013); Lippens et al. unpublished data) as well as longitudinal sampling surveys carried out in rodent communities of Senegal since the 1980s (http://www.bdrss.ird.fr/bdrsspub_form.php (Granjon and Duplantier 2009) to select sampling sites (villages or towns) into three categories of ‘*invasion status*’ for each invasion route: (i) coastal sites of long-established invasion, where rats/mice have settled in large and permanent populations since at least the 18^th^-19^th^ century and have excluded native rodents, (ii) sites at invasion front where invasive rodents have recently arrived (10-30 years ago) and currently co-occur with native ones, and (iii) non-invaded sites where only native species are known to occur. Three to six sites were systematically sampled for each category (Fig. 1). Details on rodent trapping and identification as well as autopsy procedures were provided elsewhere (Dalecky, et al. 2015; Diagne, et al. 2016). For bacterial 16S rRNA amplicon sequencing, spleens were carefully removed, placed in RNAlater storage solution, stored at 4°C one night and then at -20°C until further analyses.

**Bacterial 16S rRNA amplicon sequencing.** Data on bacterial communities considered in this study were retrieved from a 16S rRNA amplicon sequencing approach described in (Galan, et al. 2016). It enabled to get an almost complete inventory of the bacterial community infecting the rodents sampled in this study. Briefly, genomic DNA was extracted from the spleen using DNeasy 96 Tissue Kit (Qiagen) and screened via Illumina MiSeq sequencer after a PCR amplification using a slightly modified version of the universal primers of (Kozich, et al. 2013) to amplify a 251bp portion of the 16S rRNA V4 region (V4F: GTGCCAGCMGCCGCGGTAA; V4R: GGACTACHVGGGTWTCTAATCC). This hyper variable region of the 16S rRNA gene allows an accurate taxonomic assignation of bacteria to the genus level (Claesson, et al. 2010) for a wide range of bacteria. The technical procedures and bioinformatics analyses employed to get data considered in this work are presented in detail elsewhere (Galan, et al. 2016). Briefly, the MiSeq data was processed with the program MOTHUR v1.34, and taxonomic assignation was performed using the SILVA SSU Ref database v119 as a reference and refine using phylogenetic analyses and blast in GenBank. The reads generated by the Illumina MiSeq sequencing were clustered into Operational Taxonomic Units (OTUs) with a 3% divergence threshold. Finally, the data were filtered to eliminate false positive and false negative results for each bacterial OTU. The presence of a given OTU within a given rodent was next validated using systematic technical replicates. Results were then summarized in a presence/absence dataset with only bacterial OTUs known or suspected to be pathogenic for human or wildlife kept for further investigation.

**Data analyses.** Each invasion route was investigated separately. In order to avoid any confounding effects in the data analyses, we considered the factor ‘*invasion status*’ that combines the host species and site category on the invasion route. It included four categories: A) invasive species alone in sites of long-established invasion; B) invasive species at the invasion front; C) native species at the invasion front and D) native species at non-invaded sites.

To elucidate whether the composition of the bacterial communities was structured at site level by the host species and/or the invasion status, we constructed dissimilarity matrices based on Bray-Curtis (BC) β-diversity index obtained using data counts of infected individuals in each host population. The structuration of the bacterial communities was then visualised with a hierarchical clustering approach on BC dissimilarity matrices using a Ward linkage function (Murtagh and Legendre 2014). The effects of host species and invasion status were tested using a permutational multivariate analysis of variance (permanova) performed on distance matrices. These analyses were carried out with vegan v2.4.2 (Oksanen, et al.) and phyloseq v1.19.1 (McMurdie and Holmes 2013) R packages.

We next applied individual-level generalized linear mixed models (GLMMs) to test for patterns of enemy release, novel weapon, spillback, and dilution/density hypotheses. We used individual species richness (number of OTUs recorded in a single individual host) and OTUs prevalence (infection by a specific OTU for which prevalence in the dataset reached at least 5%) as response variables. When an OTU was found exclusively or mainly (more than 95% of infected hosts) in only one host species, variations in specific prevalence were tested using a dataset restricted to individuals of this host species only. We assumed a binomial distribution (quasibinomial if overdispersion) for incidence data and a Poisson distribution (negative binomial if overdispersion) for species richness data. The full model included individual host factors (*gender* and *body mass*), *invasion status* (A, B, C and D) and pairwise interactions between host factors and specific invasion status. *Sampling site* was considered as a random factor. The best-fitting model was selected based on a goodness-of-fit criterion, the Akaike’s Information Criterion with correction for samples of finite size (AICc). We then chose the most parsimonious model among those selected within two AIC units of the best model. *P*-values were obtained by stepwise model simplification using likelihood-ratio tests (Crawley 2007). For each final model, linear regression residuals were checked to ensure that regression assumptions regarding normality, independence and homogeneity of variance were met (Crawley 2007). Multiple pairwise comparisons were performed via Wilcoxon test with Holm’s method correction for *p-value* adjustment. All model analyses were performed using lme4 v1.1-8 (Bates, et al. 2015) and MuMIn v1.15.1 (Bartoń 2013) R packages.

**Ethical statements.** Trapping campaigns inside villages and private lands were conducted with permission given by the adequate institutional and familial authorities. None of the rodent species investigated in the present study has protected status (see list of the International Union for Conservation of Nature). All procedures were designed to minimize animal suffering and were approved by the official guidelines of the American Society of Mammalogists (Sikes, et al. 2011). The spleen samples were sent in dry ice to Montpellier (France) to be conserved in experimental animal room from CBGP (34-169-003). All transfer and conservation procedures were carried out paying attention to comply with current international legislations.

**Figure 4:**
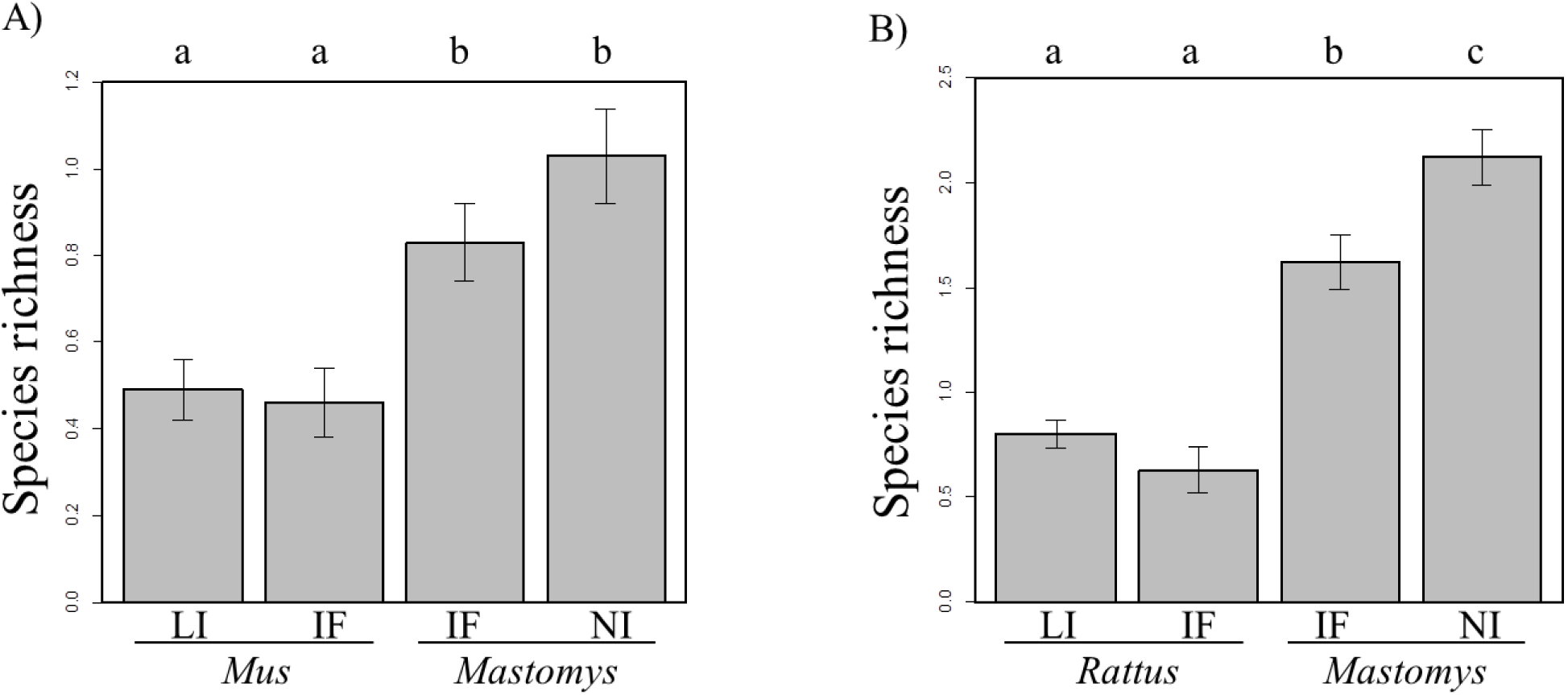
Difference in species richness (number of bacterial OTUs found in one host individual) between the specific invasion statuses from the A) mouse invasion route and B) rat invasion route. Error bars represent standard error for species richness data. Different letters above boxplots indicate significant difference between two categories of specific invasion status. Legend: *Mus*: *Mus musculus domesticus*; *Mastomys*: *Mastomys erythroleucus* for the mouse invasion route; *Ma. erythroleucus* and *Mastomys natalensis* in IF sites, *Ma. natalensis* in NI sites for the rat invasion route; LI: sites of long-established invasion; IF: invasion front; NI: non-invaded sites.

**Figure 5:**
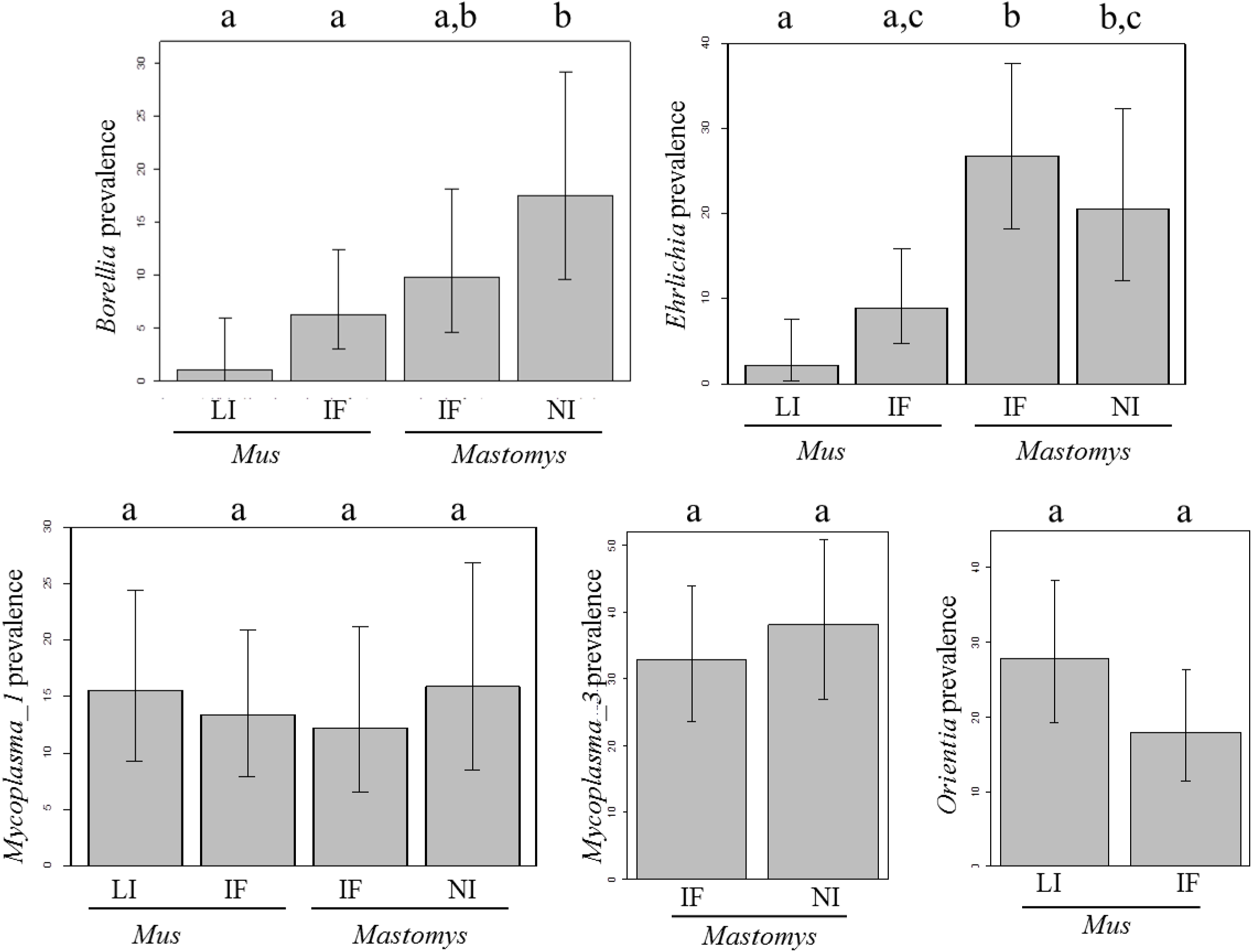
Difference in OTU prevalence between the categories of specific invasion status from the mouse invasion route. Only OTUs showing overall prevalence > 5% in the dataset are showed. Error bars represent 95% confidence intervals calculated with Sterne’s exact method for prevalence data. Different letters above boxplots indicate significant difference between two categories of specific invasion status. Legend: *Mus: Mus musculus domesticus*; *Mastomys*: *Mastomys erythroleucus*; LI: sites of long-established invasion; IF: invasion front; NI: non-invaded sites.

**Figure 6:**
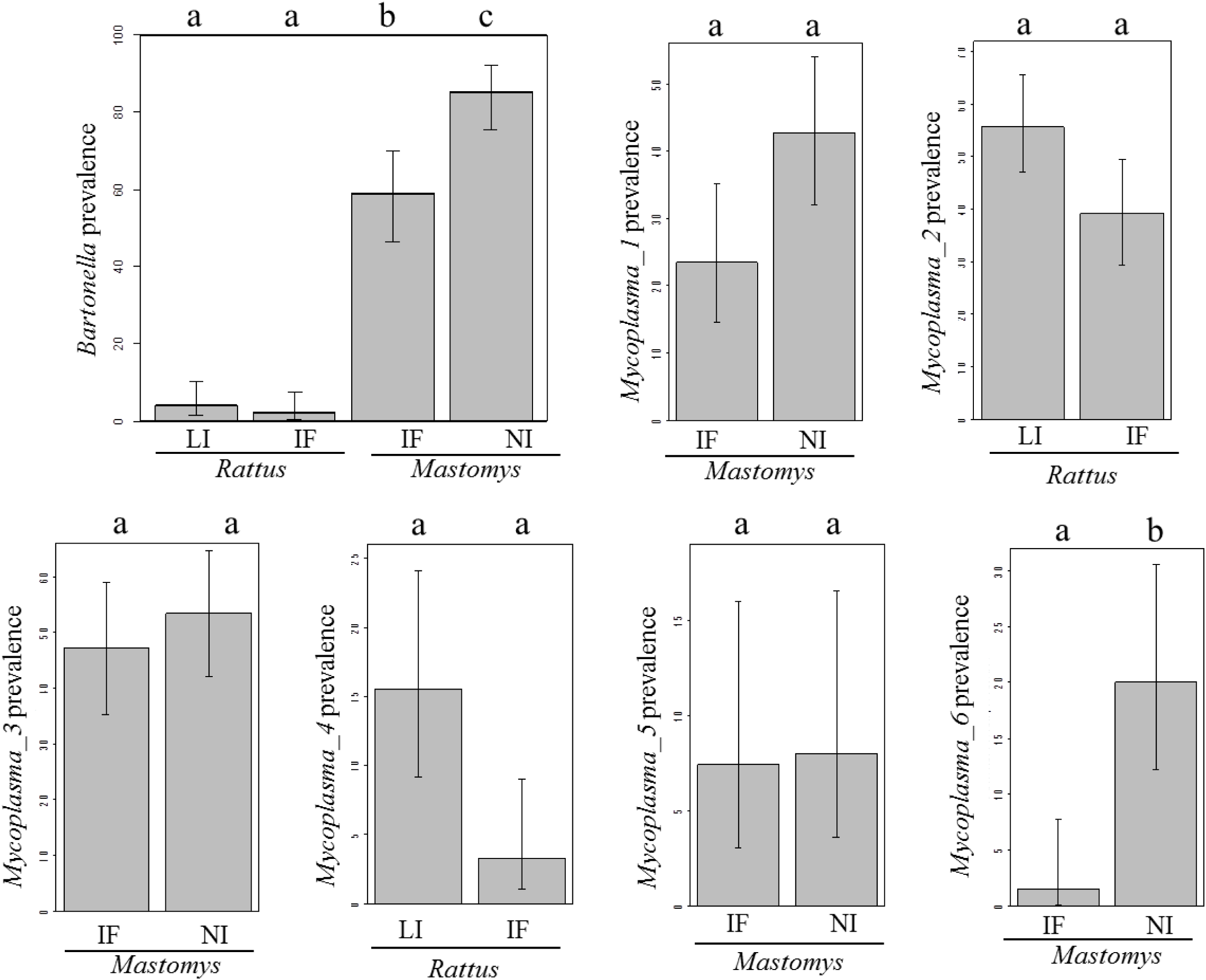
Difference in OTU prevalence between the categories of specific invasion status from the rat invasion route. Error bars represent 95% confidence intervals calculated with Sterne’s exact method for prevalence data, and standard error for species richness data. Different letters above boxplots indicate significant difference between two categories of specific invasion status. Legend: *Rattus*: *Rattus rattus*; *Mastomys*: *Mastomys* spp. (*Ma. erythroleucus* and *Ma. natalensis* in IF sites, *Ma. natalensis* in NI sites); LI: sites of long-established invasion; IF: invasion front; NI: non-invaded sites.

**Table 4:**
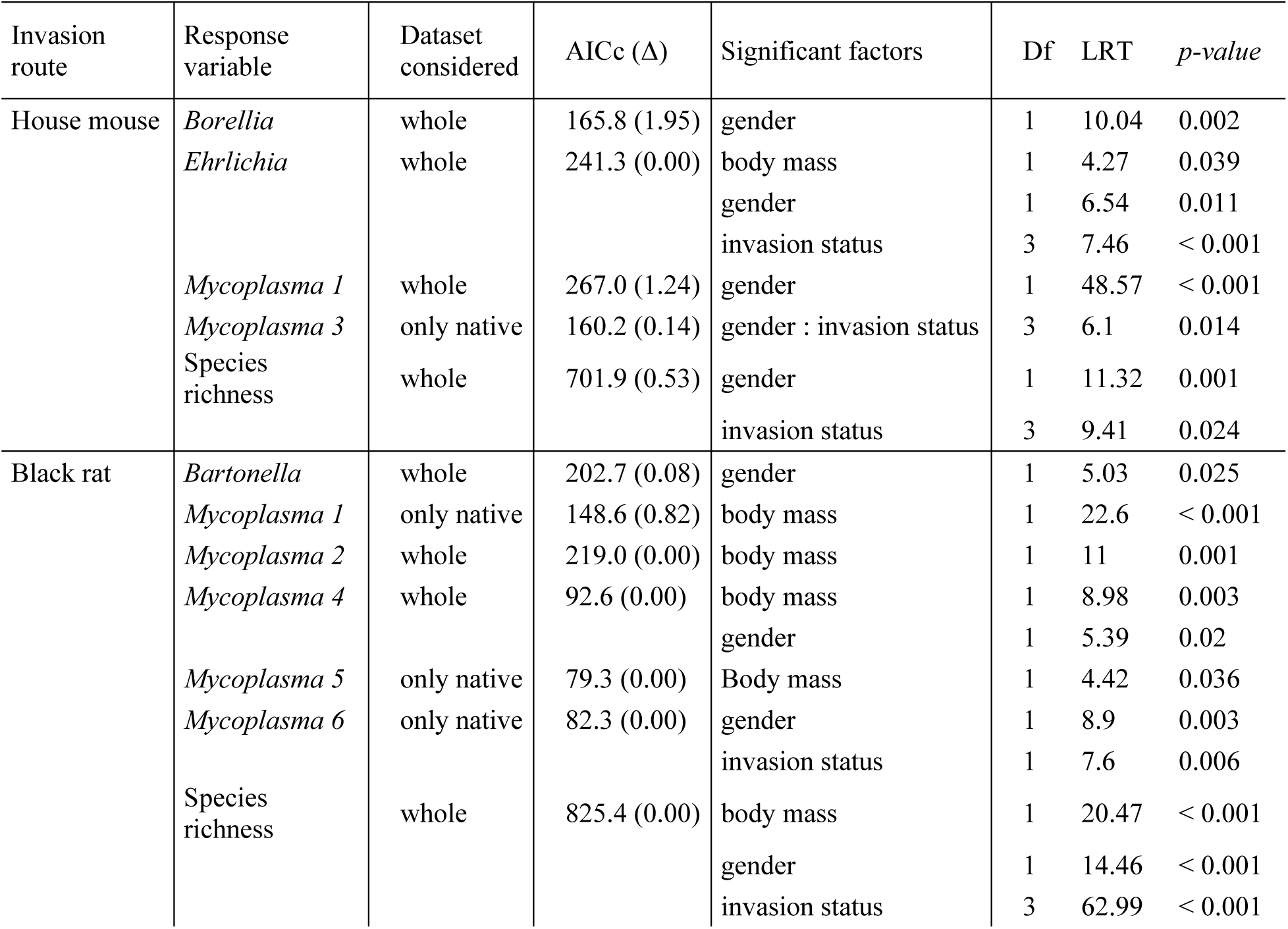
Factors shaping bacterial species richness and OTUs prevalence along both invasion routes. Significant factors are those from the most parsimonious generalized linear mixed model obtained after model selection and residuals-based validation procedures carried out for each response variable. Variations in specific prevalence were tested using a dataset restricted to a single host species when at least 95% of infected hosts by an OTU were from one species. AICc: Akaike’s information criterion with correction for finite sample size. Δ indicates the difference between the model selected and the model with the lowest AICc. M: Males; F: Females. LRT: Likelihood-ratio test value. *P-value* was considered significant when < 0.05.

## Acknowledgments

We thank Ambroise Dalecky, Khalilou Bâ, Mamadou Kane, Aliou Sow, Philippe Gauthier, Youssou Niang and Mamoudou Diallo for their participation to the field sampling in Senegal, Hélène Vignes for the MiSeq sequencing and Alexandre Dehne-Garcia for his help in the use of the CBGP HPC computational platform. We also thank Laurent Granjon, Sylvain Piry and Jean-Marc Duplantier for their precious help and advice during this work. This work was supported by the ANR ENEMI project (ANR-11-JSV7-0006). We are particularly indebted to all the people in Senegal who allowed us to carry out rodent trapping in their homes.

---

**Supplementary figure 1:**
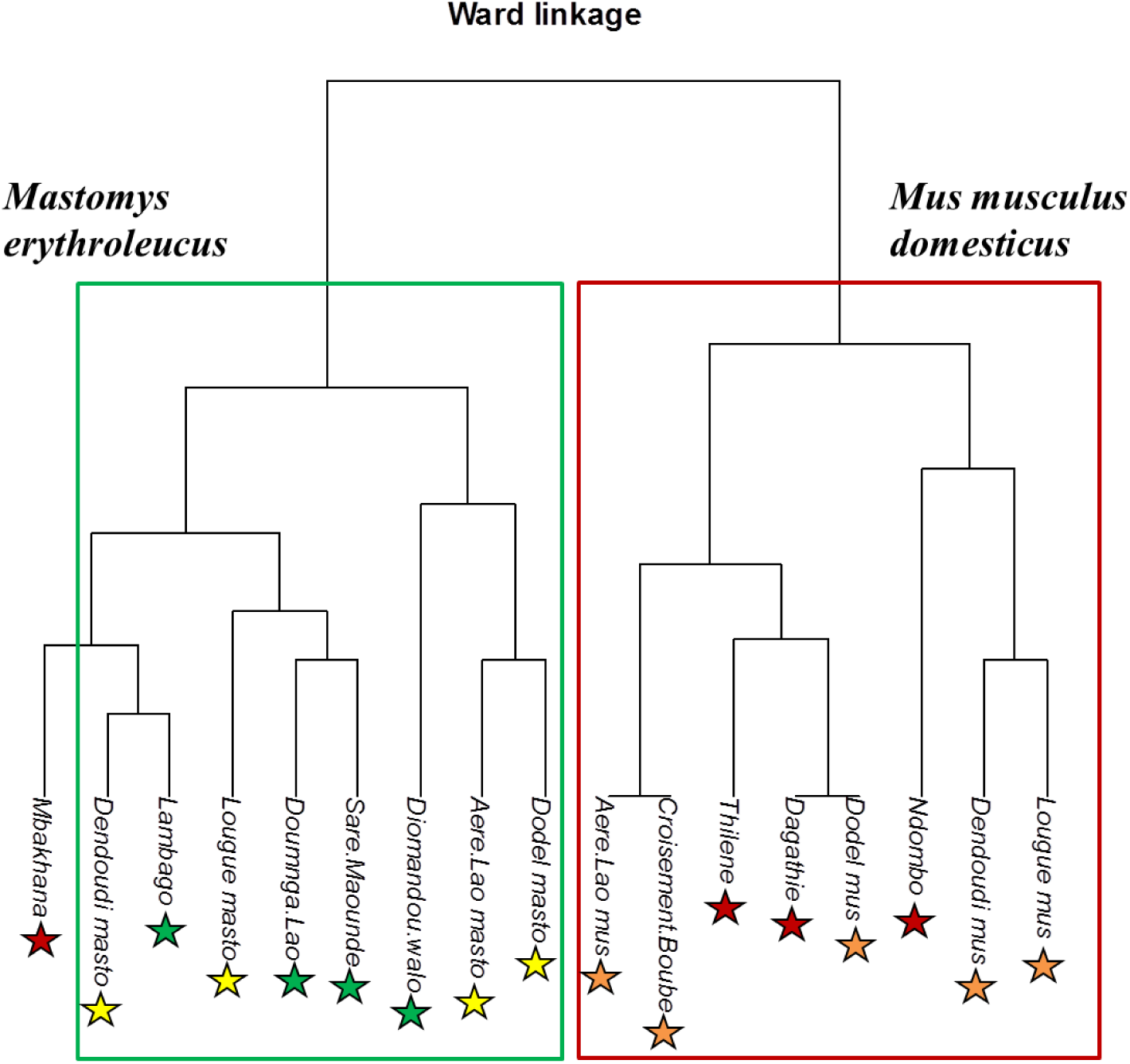
Jaccard dissimilarity index-based Ward’s hierarchical clustering of the bacterial communities described in the rodent host populations sampled along the mouse invasion route. The graph shows that the bacterial communities were clustered by the host species (permutational multivariate analyses of variance [permanova] performed on Jaccard index-based matrices: percentage of total variance explained R2 = 0.33; F = 6.82, *p = 0.001*) but not by the invasion status of sampling sites (permanova performed on Jaccard index-based matrices: R2 = 0.03; F = 0.33, *p = 0.919*). Legend: red stars: *Mus musculus domesticus* populations at sites of long-established invasion; orange stars: *M. m. domesticus* populations at invasion front sites; yellow stars: *Mastomys erythroleucus* populations at invasion front sites; green stars: *Ma. erythroleucus* populations at non-invaded sites. At invasion front sites where both native and invasive species co-occurred, “mus” or “masto” was added after the site name for distinguishing *M. m. domesticus* and *Ma. erythroleucus* populations, respectively.

**Supplementary figure 2:**
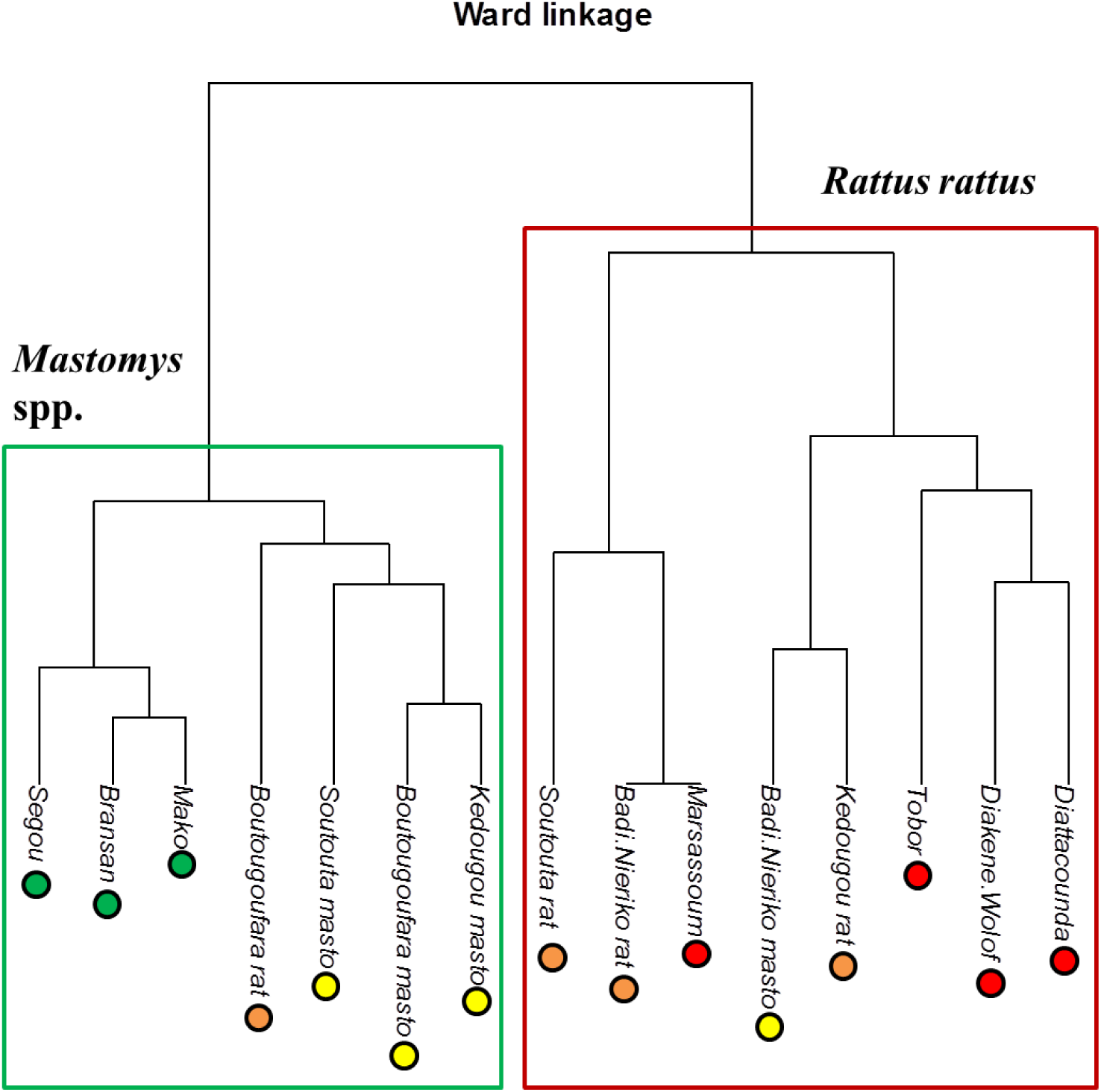
Jaccard dissimilarity index-based Ward’s hierarchical clustering of the bacterial communities described in the rodent host populations sampled along the rat invasion route. The graph shows that the bacterial communities were clustered by the host species (permutational multivariate analyses of variance [permanova] performed on Jaccard index-based matrices: percentage of total variance explained R2 = 0.39; F = 8.57, *p < 0.001*) but not by the invasion status of sampling sites (permanova performed on Jaccard index-based matrices: R2 = 0.12; F = 1.31, *p = 0.253*). Legend: red stars: *Rattus rattus* populations at sites of long-established invasion; orange stars: *R. rattus* populations at invasion front sites; yellow stars: *Mastomys* spp. populations at invasion front sites; green stars: *Mastomys* spp. populations at non-invaded sites. At invasion front sites where both native and invasive species co-occurred, “rat” or “masto” was added after the site name for distinguishing *R. rattus* and *Mastomys* spp. populations, respectively.

